# Improved binding affinity of the Omicron’s spike protein with hACE2 receptor is the key factor behind its increased virulence

**DOI:** 10.1101/2021.12.28.474338

**Authors:** Rajender Kumar, N. Arul Murugan, Vaibhav Srivastava

**Affiliations:** Division of Glycoscience, Department of Chemistry, School of Engineering Sciences in Chemistry, Biotechnology and Health, KTH, Royal Institute of Technology, AlbaNova University Center, 106 91 Stockholm, Sweden; Department of Computer Science, School of Electrical Engineering and Computer Science, KTH Royal Institute of Technology, S-100 44 Stockholm, Sweden; Department of Computational Biology, Indraprastha Institute of Information Technology, New Delhi, Delhi 110020, India

**Keywords:** SARS-CoV-2, Omicron, hACE2 receptor, Molecular dynamics simulation, Receptor-binding domain, Receptor-binding motif

## Abstract

The new variant of SARS-CoV-2, Omicron, has been quickly spreading in many countries worldwide. Compared to the original virus, Omicron is characterized by several mutations in its genomic region, including spike protein’s receptor-binding domain (RBD). We have computationally investigated the interaction between RBD of both wild-type and omicron variants with hACE2 receptor using molecular dynamics and MM-GBSA based binding free energy calculations. The mode of the interaction between Omicron’s RBD to the human ACE2 (hACE2) receptor is similar to the original SARS-CoV-2 RBD except for a few key differences. The binding free energy difference shows that the spike protein of Omicron has increased binding affinity for the hACE-2 receptor. The mutated residues in the RBD showed strong interactions with a few amino acid residues of the hACE2. More specifically, strong electrostatic interactions (salt bridges) and hydrogen bonding were observed between R493 and R498 residues of the Omicron RBD with D30/E35 and D38 residues of the hACE2, respectively. Other mutated amino acids in the Omicron RBD, e.g. S496 and H505, also exhibited hydrogen bonding with the hACE2 receptor. The pi-stacking interaction was also observed between tyrosine residues (RBD-Tyr501: hACE2-Tyr41) in the complex, which contributes majorly to binding free energies suggesting this as one of the key interactions stabilizing the complex formation. The structural insights of RBD:hACE2 complex, their binding mode information and residue wise contributions to binding free energy provide insight on the increased transmissibility of Omicron and pave the way to design and optimize novel antiviral agents.

## 1. Introduction

After its emergence in 2019, SARS-CoV-2 (Severe acute respiratory syndrome coronavirus type 2) has rapidly affected the worldwide population and caused severe pandemics [1, 2]. The four main structural proteins present in coronaviruses are spike (S), envelope (E), nucleocapsid (N), and membrane (M) proteins [2, 3]. The primary function of the S protein is to bind to the receptor Angiotensin-Converting Enzyme 2 (ACE2) that helps the virus enter the host cell [4]. Thus, the S protein plays a critical role in the initial stage of infections for disease-causing coronaviruses. Therefore, all studies focused on understanding the detailed mechanism of S protein-ACE2 interaction is crucial. This protein-protein complex is considered a prime target for developing new drugs and vaccines.

Several SARS-CoV-2 variants have been reported so far around the world. Based on the evidence of their increased transmissibility, disease severity and/or possible reduced vaccine efficacy, four variants have been universally categorized as variants of concern (VOCs) (https://www.ecdc.europa.eu/en/covid-19/variants-concern). A new SARS-CoV-2 variant (belonging to the Pango lineage B.1.1.529) was identified in South Africa at the end of November 2021[5]. The World Health Organization (WHO) named this variant Omicron and classified it as a VOC. Based on the changes that occurred at the sequence level in the Omicron variant, it is assumed that it may be more transmissible than the original SARS-CoV-2 virus and other variants (e.g. alpha, delta etc.); however, no in-depth study based on experimental and computational methods has supported this assumption. Nevertheless, the number of SARS-CoV-2 infected cases has increased worldwide and poses a major threat to the world healthcare system. Recently, researchers from the University of Hong Kong explained the reason behind the increased transmissibility of the Omicron than previous SARS-CoV-2 variants [6]. They showed that Omicron infects and multiplies 70 times faster than the original virus and Delta variant and in the human bronchus. Another study by Grabowski et al. (2021) reported the exponential growth of the Omicron over the four weeks (November 8– December 5, 2021) with an estimated doubling time of 3.38 days [7]. Indeed, WHO has also recognized that Omicron is spreading at an exceptional rate.

It was found that compared to the original wild type (wt) virus, the Omicron variant is characterized by 30 amino acid mutations, three small deletions and one small insertion in the spike protein (https://www.who.int). Apparently, 15 of these mutations are located in the re-ceptor-binding domain (RBD). Out of these, 10 mutations are specific to the receptor-binding motif (RBM) (Figure 1), through which the viral spike protein interacts directly with the angiotensin-converting enzyme 2 (ACE2) receptors. In addition, the Omicron variant carries some changes and deletions in other genomic regions (https://www.ecdc.europa.eu/). The initial step of the infection process is binding the spike RBD to the human ACE-2 receptor, indicating that viral transmissibility and virulence largely depend on the interaction between these proteins. Mutations in the RBD of the spike protein or other surface structures can change the new variant’s antigenic property, reducing the neutralization activity of antibodies and resulting in a higher risk of reinfection or decreasing vaccines effectiveness.

**Figure 1.**
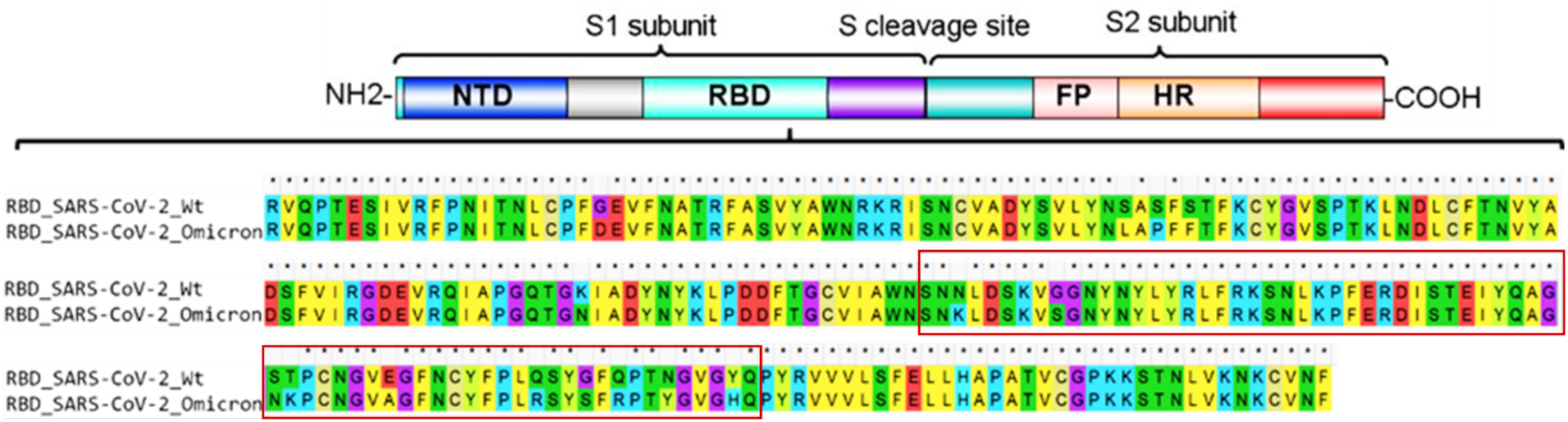
The Coronavirus spike protein domain architecture, and sequence alignment of the original RBD of SARS-CoV-2 and RBD of the SARS-CoV-2 Omicron variant. Asterisk denotes identical sequences. The red rectangular box indicates the receptor-binding motif of RBD. NTD, N-terminal domain; RBD, receptor-binding domain; FP, fusion peptide; HR, heptad repeat.

Therefore, it is urgently needed to study the mutation pattern of the Omicron SARS-CoV-2 and the mechanism of virulence or pathogenesis to develop effective measures. In this study, we have investigated how different mutations in the receptor-binding domain (RBD) of the spike protein of the Omicron variant affect its binding to the ACE2 receptor. This fundamental understanding of sequence to structural level interaction between the Omicron spike RBD and hACE2 is highly demanded in developing new treatments for coronavirus infections. We have employed force-field molecular dynamics and subsequently the implicit solvent binding free energy calculations to analyze the relation binding strength of Omicron variant when compared to wild type with hACE2 receptor.

## 2. Result and discussion

The SARS-CoV-2 surface spike protein interacts with the human ACE2 receptor through its receptor-binding domain (RBD) and facilitates the entry of the virus into the host cell. The structural information of many SARS-CoV-2 RBD-hACE2 complexes have been reported in the PDB database (https://www.rcsb.org/) that provides in-depth information about their binding interface. A receptor-binding motif (RBM, about 69 amino acids) of the RBD directly contacts and binds to the ACE2 receptor, clearly indicating that any mutation in RBM might play a vital role in the infection process [8]. Indeed, many naturally occurring mutations in SARS-CoV-2, mainly those in the RBD, affected its binding to the ACE2 receptor [9]. Out of about 30 amino acid substitutions reported in the Omicron spike protein, many are present in RBD/RBM (https://www.who.int/). Here, we performed a comparative sequence analysis to examine the key residues changed over time within the RBM of all reported SARS-CoV-2 (Figure 2). It was found that in total 22 different positions (Figure 2) were substituted with various amino acids in the RBM of SARS-CoV-2 variants including the Omicron. In particular, the Omicron RBM has a total of 10 mutations: N440K, G446S, S477N, T478K, E484A, Q493R, G496S, Q498R, N501Y, and Y505H (Figure 2). Interestingly, the residues E484 and N501 (as in wt-SARS-COV-2) were substituted many times by different amino acids in different variants (Figure 2). In case of Omicron, these amino acids were mutated with alanine and tyrosine (E484A and N501Y). In addition, six mutations at positions G446S, E484A, Q493R, G496S, Q498R, and Y505H were only observed in Omicron, while rest of the four mutations were also present in other variants (Figure 2). The six unique mutations mentioned above in the Omicron RBM may directly affect the binding of the spike protein to the host cell ACE2 receptor.

**Figure 2.**
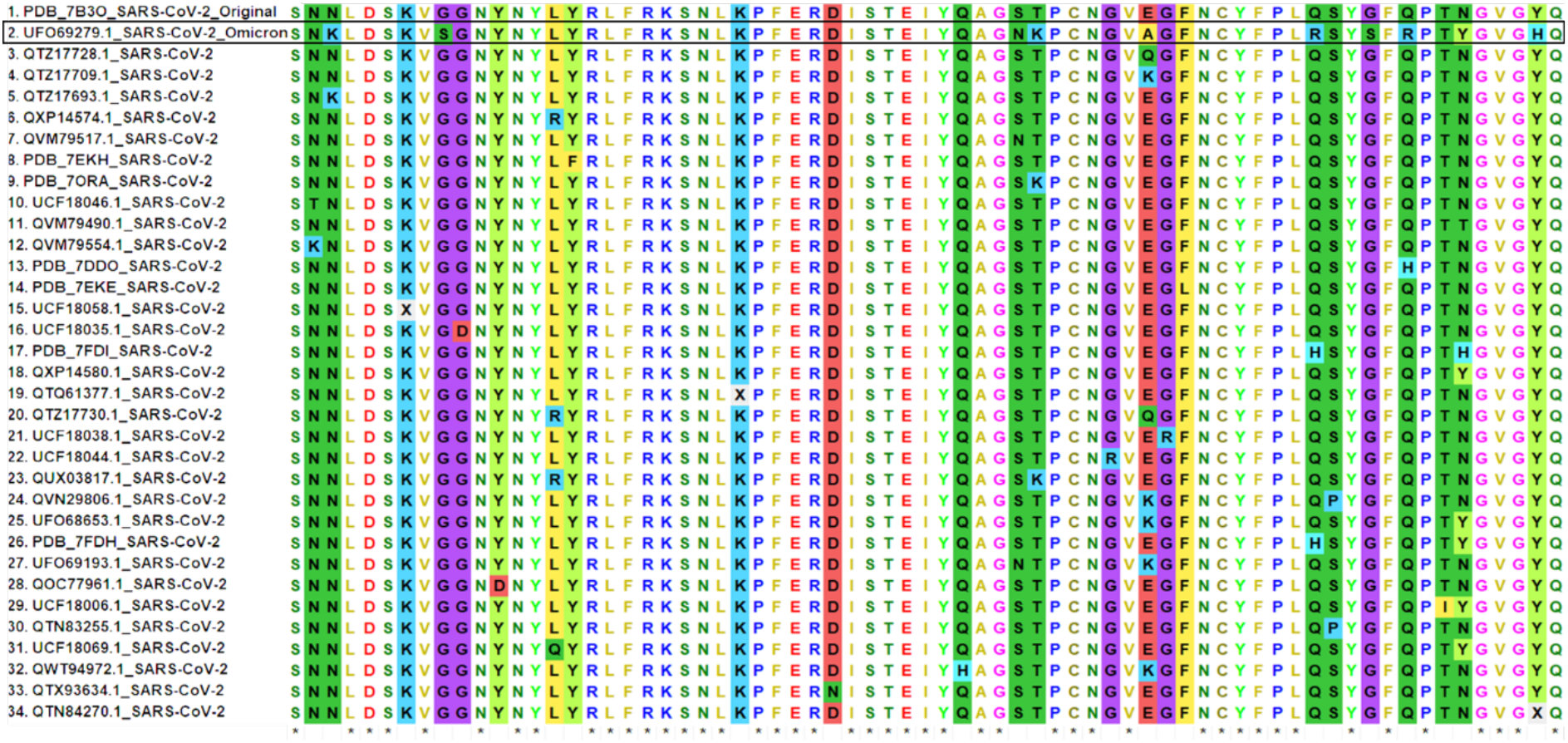
Comparative sequences analysis of the spike RBM of SARS-CoV-2. A total of 34 representative sequences were aligned. Asterisk denotes identical sequences.

As recently reported by Niu et al., (2020), the RBM Q498H substitution found in pangolin CoVs, enhanced its binding capacity to ACE2 receptor homologous of mouse, rat and European hedgehog [10]. The glutamine residue at this position is replaced by arginine in the Omicron variant (Figure 2); however, the effect of this crucial mutation has not been studied so far computationally. This mutation has been shown to make hydrogen bonds and salt bridges with ACE2 (the residues involved are Q42 and D36) in a very recent study based on cryo-electron microscopy [11]. A similar type of study, as earlier mentioned [10], can also be interesting to investigate the effect of this mutation on ACE2 homologous in other hosts. In order to better understand the structural insights of the Omicron RBD binding mode to ACE2 receptor, we have performed a computational modelling and large-scale molecular dynamics simulation studies.

The classical molecular dynamic simulation studies were carried out to explore the protein-protein interaction between the Omicron RBD and the hACE2 receptor, and the results were compared to wild type RBD-hACE2 complex. The mutated and the wild type complexed structures were analysed after 100 ns MD simulation. We observed that an overall protein complex was stable except for some flexibility in loops regions during the MD simulation. The computed root mean square fluctuation (RMSF) which is proportional to the thermal factor (Figure S1a) shows that the conformational flexibility is not altered significantly in the case of the Omicron variant when compared to the wild type. The same is the case for hACE2 receptor in these two complexes (Figure S1b. The binding free energies computed for the spike RBD with the hACE2 receptor for the wild type and the Omicron variant are listed in the Table 1. As can be seen, the binding free energies for the Omicron-spike with hACE2 receptor is lower than that of wt-spike by −8.6 kcal/mol. This clearly indicates increased binding affinity towards the hACE2 receptor for the Omicron variant associated spike protein, which can be directly associated with its increased infection rate. Further observations (Table I) suggest that the major driving force for the increased binding affinity can be attributed to the increased electrostatic and hydrophobic interactions. Indeed, in the case of the Omicron spike the electrostatic interaction energy difference was almost double when compared to the wt-spike. However, the increased binding potential due to electrostatic contribution is largely compensated by the polar solvation free energies, which is comparably larger for the Omicron variant. The sum of electrostatic and polar solvation energy is, in fact, more positive for the Omicron variant suggesting that the stabilization is largely due to hydrophobic interactions. As can be seen from Table 1, the difference in van der Waals interactions was −15.6 kcal/mol (Table I) between the Omicron and wild type. We have also carried out the residue wise decomposition analysis of binding free energies, and the results are presented in figure 3.

**Figure 3:**
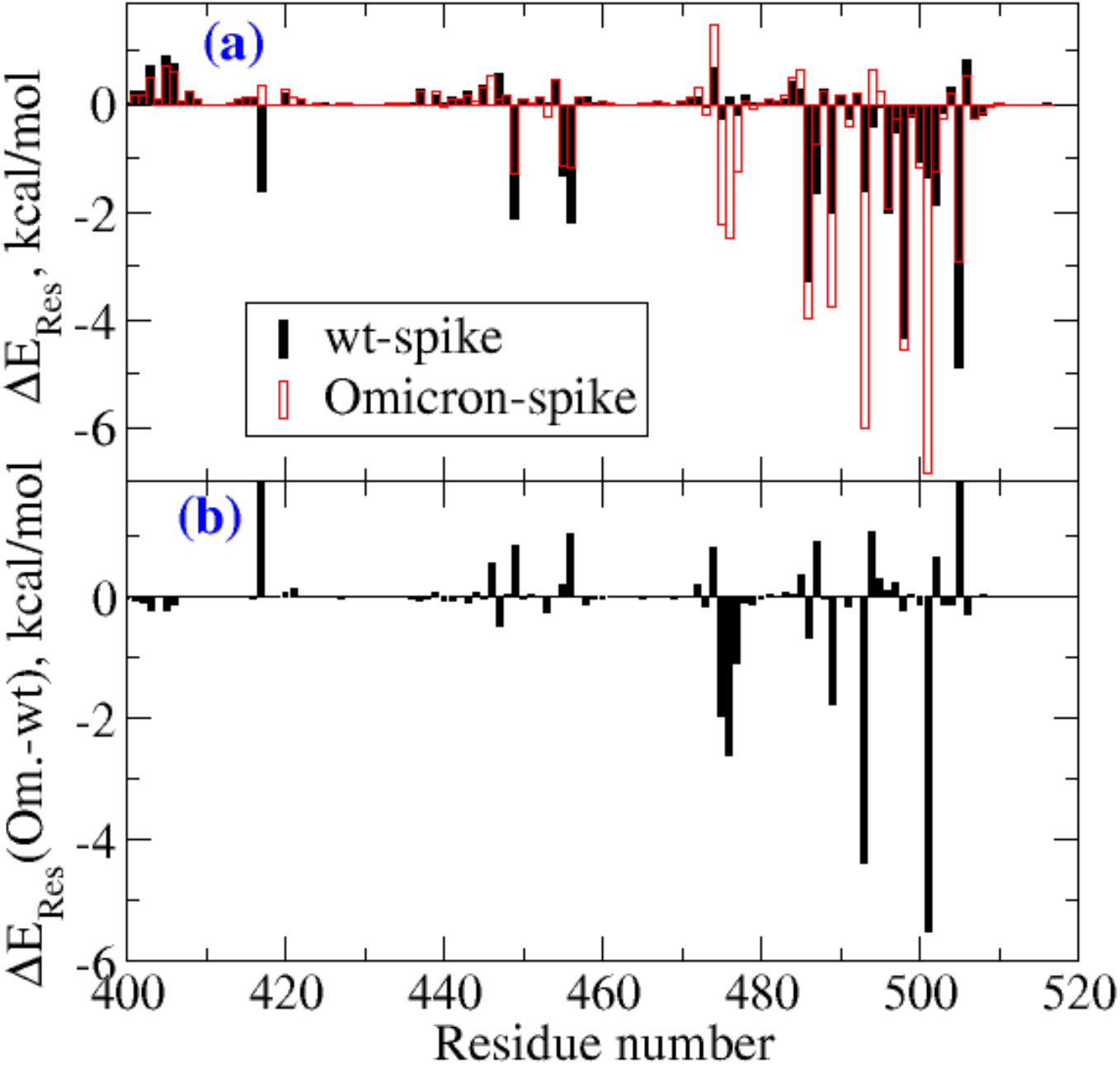
(a) Residue-wise contributions for the binding free energies. The results are shown only for residues in RBD of the spike protein. Further, the numbering of residues is shifted by three for the Omicron to account for the three deletions. (b) The difference in residue-wise contributions to binding free energies between spike proteins of the Omicron variant and the wild type (wt)

**Table I.** Binding free energies of the Omicron spike RBD and hACE-2 complex.

| System | $\Delta E_{\text{vdw}}$ | $\Delta E_{\text{elec}}$ | $\Delta E_{\text{psol}}$ | $\Delta E_{\text{npsol}}$ | $\Delta G$, kcal/mol |
| --- | --- | --- | --- | --- | --- |
| Wild type | -85.28 | -856.42 | 922.17 | -12.91 | -32.43 |
| B11529 | -100.84 | -1714.73 | 1787.97 | -13.39 | -41.00 |

The subplot 3a shows the residue wise decomposition of binding free energies for the RBD domain of the wild type and Omicron variant, while the subplot 3b shows the difference in residue-wise binding free energies between the Omicron variant and the wild type. A positive sign refers to destabilizing residues in the case of the Omicron, while the negative sign shows the stabilizing residue wise contribution. The analysis shows that 475-477, 489, 493, and 501 residues contributed to the increased binding potential of the Omicron variant. All the residues contribute to binding free energies lower than −1.0 kcal/mol. In particular, the residues 476, 493 and 501 contribute with −2.613, −4.385 and −5.49 kcal/mol. As we will see, the major contributions are due to hydrophobic, electrostatic and hydrogen bonding interactions. Further, to understand the increased binding affinity, residue level contributions, hydrogen bonding, and salt-bridges were investigated between the protein-protein complexes.

In the Omicron-RBD–hACE2 complex, a total of 11 significant hydrogen bonds and three different salt bridge ion pairs were observed. The four key hydrogen bonds, T500-D355, G502-K353, R493-D30 and R498-D38 were found with occupancies of 54.20%, 48.60%, 39.20% and 28.80%, respectively (Table II). In the wt-RBD–hACE2 complex, a total of 14 hydrogen bonding interactions and one salt bridge ion pair were observed. We also observed that some hydrogen bonding interactions are similar in both complexes with different hydrogen bond occupancies. In the wt-RBD–hACE2 complex, the highest hydrogen bond occupancy, 70.50%, was observed for the Y449-D38 pair, while the same pair in Omicron was only 17.10% (Table II). This lower occupancy is because in the Omicron RBD:hACE2 complex, D38 of hACE2 is also occupied with two residues of the Omicron RBD, the mutated R498 via salt bridge interaction and another mutated S496 via hydrogen bond. The hydrogen bond between G502-K353 was found with occupancy 48.60% in both wild type and Omicron complexes.

**Table II.** Hydrogen bonds and their occupancies, and salt bridge ion-pairs found between spike RBM and hACE2 receptor during the entire MD simulation. The mutated residues in the Omicron are shown in bold).

| No. | Wild type-RBM: hACE2 | H-Bond<br>Occupancy<br>(%) | Omicron-RBM:<br>hACE2 | H-Bond<br>Occupancy<br>(%) | Salt bridges<br>(wild type-RBM- - hACE2) | Salt bridges<br>(Omicron-RBM- -<br>hACE2) |
| --- | --- | --- | --- | --- | --- | --- |
| 1 | Tyr449-Asp38 | 70.50 | Try449-Asp38 | 17.10 | Lys417- -Asp30 | **Arg493-** -Asp30 |
| 2 | Thr500-Asp355 | 38.10 | Thr500- Asp355 | 54.20 |  | **Arg493-** -Glu35 |
| 3 | Thr500-Tyr41 | 18.40 | - | - |  | **Arg498-** -Asp38 |
| 4 | Gln493- Glu35 | 38.20 | **Arg493-** Asp30 | 39.20 |  |  |
| 5 | - | - | **Arg493-** Glu35 | 11.10 |  |  |
| 6 | Gln498- Asp38 | 26.90 | - | - |  |  |
| 7 | Gln498- Lys353 | 20.80 | **Arg498-** Asp38 | 28.80 |  |  |
| 8 | Gly502- Lys353 | 48.60 | Gly502 – Lys353 | 48.60 |  |  |
| 9 | Tyr505- Glu37 | 22.70 | **His505-** Ala386 | 16.50 |  |  |
| 10 | Gly496- Lys353 | 14.00 | **Ser496-** Asp38 | 16.10 |  |  |
| 11 | Ser494- His34 | 14.90 | Ser494- His34 | 12.20 |  |  |
| 12 | Asn487- Tyr83 | 28.30 | Asn487- Tyr83 | 9.10 |  |  |
| 13 | Ala475- Gln24 | 11.10 |  |  |  |  |
| 14 | Lys417- Asp30 | 33.50 | - | - |  |  |
| 15 |  | - | Asn487 – Gln24 | 13.90 |  |  |
| 16 | Gly446-Gln42 | 9.50 | - | - |  |  |

A key interacting interface residue glutamine, present at two different positions, 493 and 498, in the wild type was mutated to positively charged amino acid arginine in the variant Omicron (Figure 2, Table II). Both these mutated residues play an important role in binding interaction of RBD–hACE2 complex not only via hydrogen bonds but also through salt bridges (Figure 4). It is worth mentioning that these interactions were consistently observed during the entire MD simulations. These salt bridges did not occur in the wild type RBD:hACE2 complex mainly because of glutamine residues at these positions. Other hydrogen bond-forming residues in the RBD–hACE2 complex were found with lower occupancies, and some of them are mutated in the Omicron; for example, residues pairs H505-A386, and S496-D38. Altogether, the mutated residues resulted in a stronger interaction between the Omicron RBD and hACE2 via hydrogen bonds and salt bridges.

**Figure 4.**
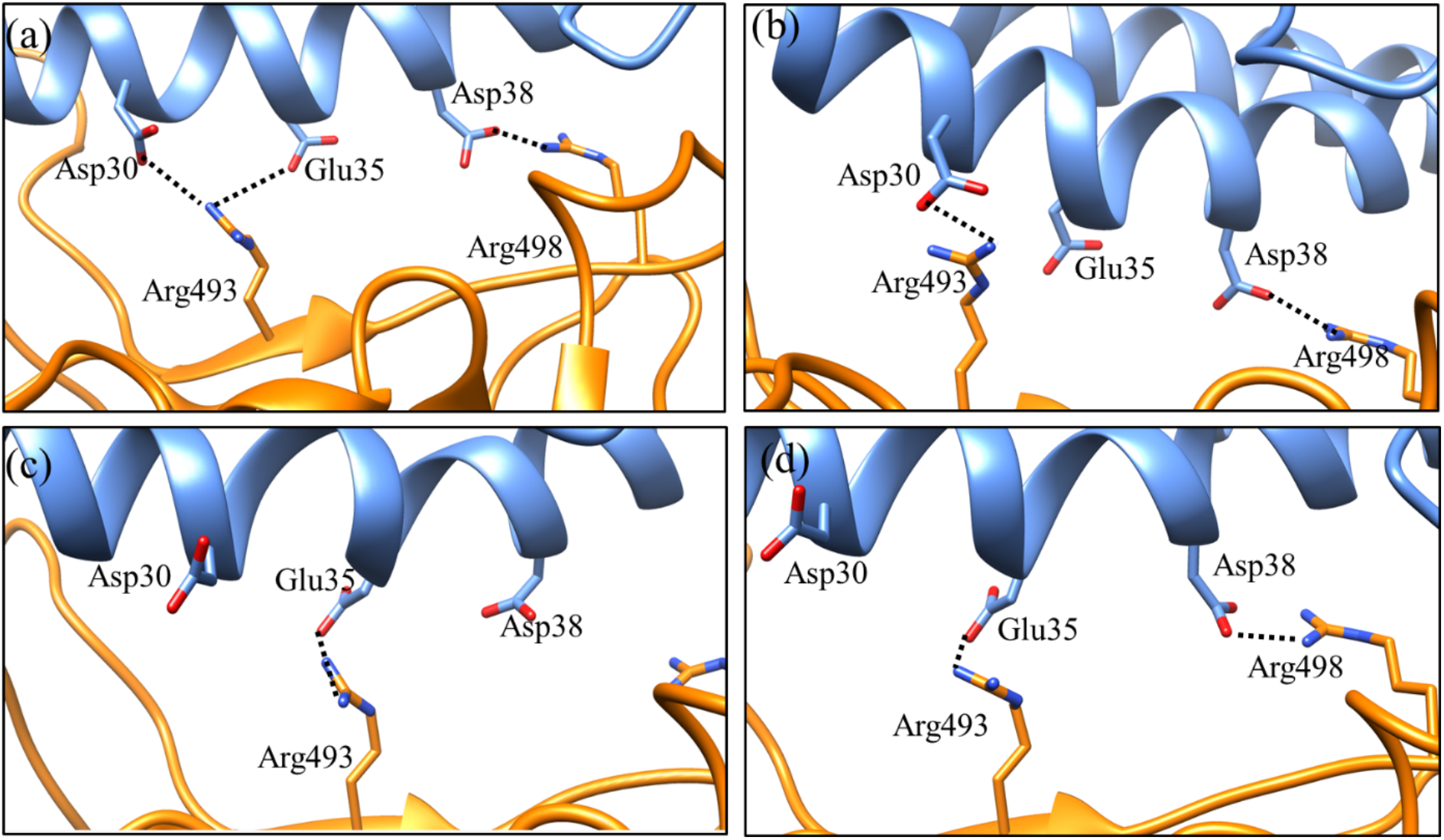
The electrostatic interaction involving residues and their interactions observed during MD simulation, (a) all possible salt bridge pairs Arg493 -- Asp30, Arg493 -- Glu35 and Arg498 -- Asp38 observed during entire 100 ns MD simulation, (b) salt bridge pairs, Arg493 -- Asp30 and Arg498 -- Asp38, observed between 0-70 ns MD simulation, (c) salt bridge pair, Arg493 -- Glu35 observed between 80-100 ns MD simulation (d) showed salt bridge pairs Arg493 – Glu35 and Arg498 -- Asp38 observed for a very short time between 80-85 ns MD simulation. The hACE2 receptor is shown in blue colour and the spike RBD by orange.

It has been reported that in the original SARS-CoV-2, the residue K417 forms salt bridge, and hydrogen bond with hACE2 D30 [1]. However, in the Omicron the K417 is mutated to asparagine residue resulting in the loss of electrostatic interaction with Asp30 of the hACE2 receptor. The same mutation (K417N) in the delta variant of SARS-CoV-2 was associated with slight decrease in the ACE2-binding affinity [12]. The detailed information about the hydrogen bonding and their occupancies and electrostatic interaction ions pairs are given in Table II.

Interestingly, when comparing the salt bridges interactions in both complexes, some key differences were found. In the wt-RBD–hACE2 complex, only one salt bridge between Lys417-NH3^+^ -- ^-^OOC-Asp30 was consistently formed with an average distance of approximately 2.9Å during the MD simulation (Figure 5). Only in a few conformations, the ion pair Arg403-NHC(NH2)2_+_ -- ^-^OOC-Glu37 comes closer, as seen in the figure 5, and after a while, it showed a larger distance more than 7.0 Å. For the wt-RBD–hACE2 complex, we considered only one salt bridge, K417-D30, that is also reported in the crystal structure [1]. While in the case of the Omicron-RBD–hACE2 complex, K417 is mutated with glutamine that did not interact with hACE2 residues. The mutated residues Q493R and Q498R have a large side chain with positively charged functional groups showed electrostatic interactions with negatively charged amino acids D30, E35 and D38 of hACE2 receptor as mentioned earlier (Figure 4, Table II).

**Figure 5.**
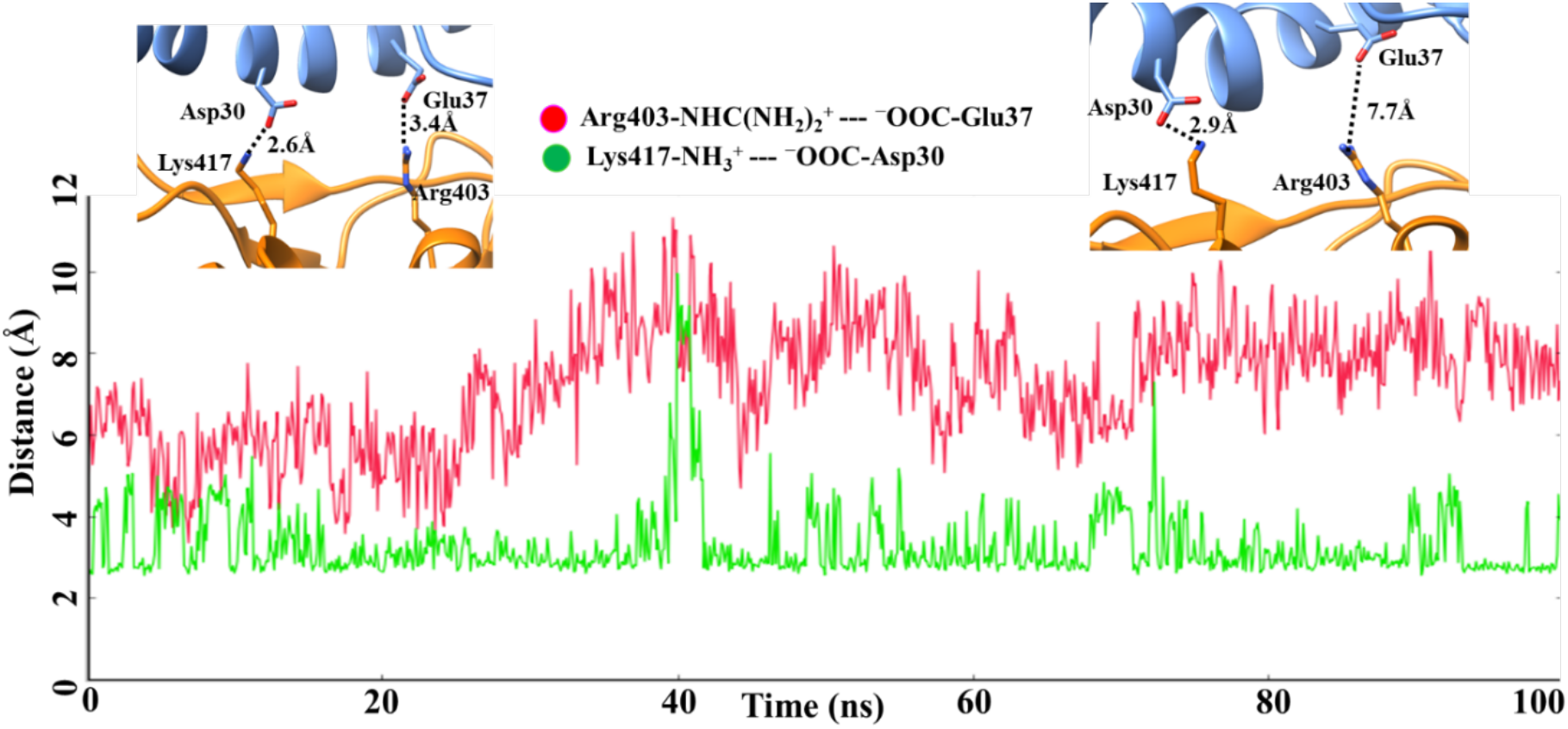
Salt bridges interactions and their corresponding distances between the wild-type spike RBD and the hACE2 receptor complex during 100ns MD simulation

Interestingly, we have observed two salt bridges consistently formed during the MD simulation via ion pair Arg493-NHC(NH2)2^+^ -- ^-^OOC-Asp30/-Glu35 and Arg498-NHC(NH2)2^+^ -- ^-^OOC-Asp38 (Figure 4). As shown in the figure 6, salt bridge pairs R493 -- D30 and R498 −D38 observed between 0-70 ns MD simulation while salt bridge pairs R493 -- E35 (average distance less than 2Å) was observed between 80-100 ns MD simulation (Figure 6). The side chain of residue R493 showed two different conformations in which one was observed near D30 for longer time while alternate conformation that comes closer to E35 for short times during MD simulations. A very short lifetime salt bridge pairs R493 – E35 and R498 -- D38 simultaneously observed between 80-85 ns MD simulation (Figure 6).

**Figure 6.**
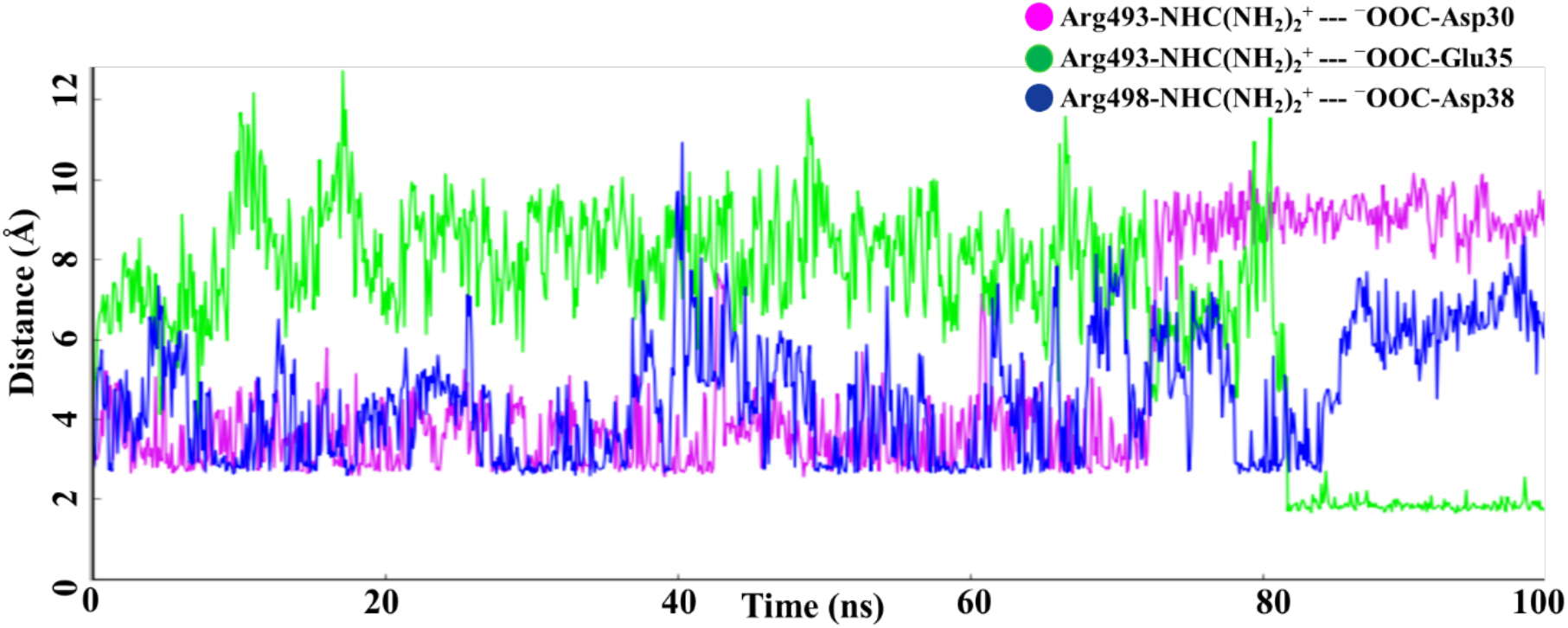
The distance between residues participating in the salt bridge (electrostatic interaction) during the entire MD simulation

Additionally, a non-canonical π–π stacking interaction was found at the interface of the Omicron RBD-hACE2 complex between two tyrosine residues located on the protein surface. The aromatic ring of the mutated Y501 residue in the Omicron RBD showed pi-stacking interaction with the aromatic ring of Y41 of the hACE2. It was found that the geometry of this interaction was T-shape, where the aromatic rings of Y41 and Y501 were perpendicular to each other. It has been earlier shown that such Tyr-Tyr interaction occurs mainly at the protein surface [13]. The average distance obtained in most of the conformations was below 4Å between aromatic rings. As McGaughey (1998) analysed earlier, the aromatic amino acids side chains can stabilise interactions in protein with a larger distance than average van der Waals radii [14]. The residue wise decomposition analysis of binding free energies showed that in fact the residue 501 of the spike protein of Omicron variant is the one dominantly contributing with as much as −5.5 kcal/mol. The mutation N501Y in the RBD reported for SARS-CoV-2 variants has been shown to increase higher ACE2 binding affinity due to improved π-π stacking [11, 15] and is also associated with increased infection and transmission [16]. In the Omicron-RBD:hACE2 complex, both tyrosine residues are located on the protein’s surface. This interaction is absent in the wild type hACE2 complex as the amino acid present at the position 501 in wild type RBD is asparagine that is mutated to tyrosine in the Omicron.

## 3. Conclusions

The rapidly spreading B.1.1.529 lineage (Omicron) has many mutations, indicating an increased risk of reinfection and tendency to escape to therapeutic vaccines and antibodies in the coming future. The RBM sequence analysis and protein-protein interaction studies on RBD and hACE2 complex using molecular dynamics and binding free energy calculations provided the detailed molecular level binding interaction pattern. Sequence and structural level investigation of the RBD reveals that the mutated residual composition in the variant Omicron exhibited an increase in the number of charged amino acids such as arginine, lysine, and aspartic acid, that contribute to electrostatic interaction in protein. Our investigation indicates the mutated RBM interface bound tightly to ACE2 in the context of binding free energies. We have observed extra salt bridges between the RBM-R493--D30-ACE2 and RBM-R493--E35-ACE2 and RBM-R493--D38-ACE2 in the Omicron variant. Interestingly, lysine residue at position 417 in the RBD of the original SARS-CoV-2, which forms a salt-bridge with D30 of hACE2, did not show any interaction with this receptor after being mutated to glutamine in Omicron. Additionally, RBM’s mutated resides formed some additional hydrogen bonding and pi-stacking interactions, which could further enhance hACE2 binding. This information could be more informative for viral communities’ scientists engaged in finding the better therapeutics for SARS-CoV-2, especially for the new variants.

## 4. Materials and Methods

### 4.1. Sequence analysis and structure modelling

For comparative analysis, sequences of the receptor-binding motif (RBM) from all reported SARS-CoV-2 variants were retrieved from NCBI database using BlastP program. Further, sequence redundancy was removed at 100% sequence identity, and the remaining representative sequences were aligned using EBI-MUSCLE program (https://www.ebi.ac.uk/Tools/msa/muscle/). For computational modelling, all 15 RBD substitutions G339D, S371L, S373P, S375F, K417N, N440K, G446S, S477N, T478K, E484A, Q493R, G496S, Q498R, N501Y, and Y505H were incorporated into the original resolved crystal structure (PDB ID: 7A91) [17] using the Mutagenesis module of PyMOL software (http://www.pymol.org/pymo) to get the Omicron spike RBD model structure. The mutated structure was energy minimized for further refinement and used for molecular dynamics (MD) simulation and binding free energy studies.

### 4.2 MD Simulation and MM-GBSA free energy calculations

Molecular dynamics simulations of the protein complexes, (i) Omicron-RBD:hACE2, and (ii) wt-RBD:hACE2, were carried out using the AMBER 18.0 package [18, 19] where f14SB force field [20] parameters were applied for proteins. The complexes were solvated with a sufficient number of water molecules (TIP3P force field has been used to describe water) and neutralized with counter ions in an orthorhombic simulation box [21]. In particular, the spike RBD-hACE2 complex is embedded in a simulation box with approximately 37000 water molecules.

The solvated systems were energy minimized and followed by this, low temperature simulations (at 30 K and 1atm pressure) were carried out. Each system was then equilibrated to a free simulation for a short time. Finally, MD simulations in ambient conditions (300 K and 1atm) were carried out. The time step for solving Newton’s equation of motion was set to 2fs. The time scale for the production runs for each complex was set to 100 ns and the trajectories were recorded every 50ps time interval. Further, the trajectories were used for computing various properties like RMSF, the radius of gyration (Rg), etc. The RMSFs and Rg results computed for the spike protein and the hACE2 receptor are shown in Figures S1 and S2, respectively. The intermolecular non-covalent interactions such as hydrogen bonding, salt bridges, and pi-stacking were analyzed using VMD [22] and UCSF Chimera software [23]. The binding free energies were computed as an average of over 2500 configurations corresponding to the last 10 ns of the production run using molecular mechanics-Generalized Born surface area approach (MM-GBSA) [24, 25].

## Author Contributions

Conceptualization, V.S and N.A.M.; methodology, R. K. and N.A.M.; software, R. K. and N.A.M.; validation, R. K. and N.A.M.; formal analysis, V.S., R. K. and N.A.M.; investigation, V.S., R. K. and N.A.M.; resources; writing-original draft preparation, review and editing, R. K., N.A.M. and V.S.; and supervision, N.A.M and V.S. All authors have read and agreed to the published version of the manuscript.

## Institutional Review Board Statement

Not applicable.

## Informed Consent Statement

Not applicable.

## Data Availability Statement

Data sharing not applicable.

## Acknowledgments

The authors would like to acknowledge the Swedish Infrastructure Committee (SNIC) resources for the projects SNIC 2021/5-119, SNIC 2021/5-1, SNIC 2021/5-39, and SNIC 2021/5-471.

## Supporting information

**Figure S1:**
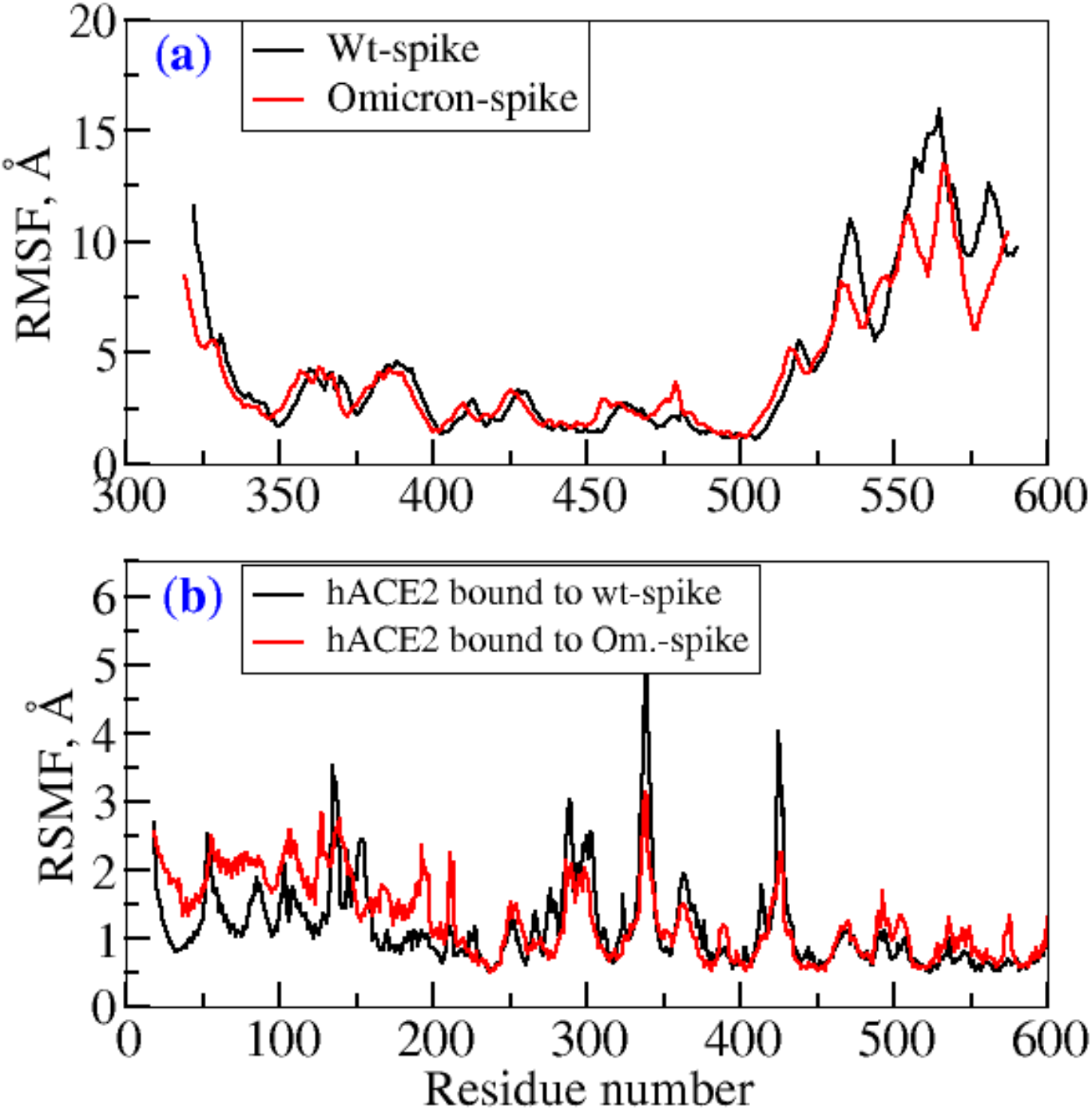
RMSFs computed for (a) spike protein and (b) hACE2 in the two complexes wt-spike: hACE2 and Omicron-spike: hACE-2

**Figure S2:**
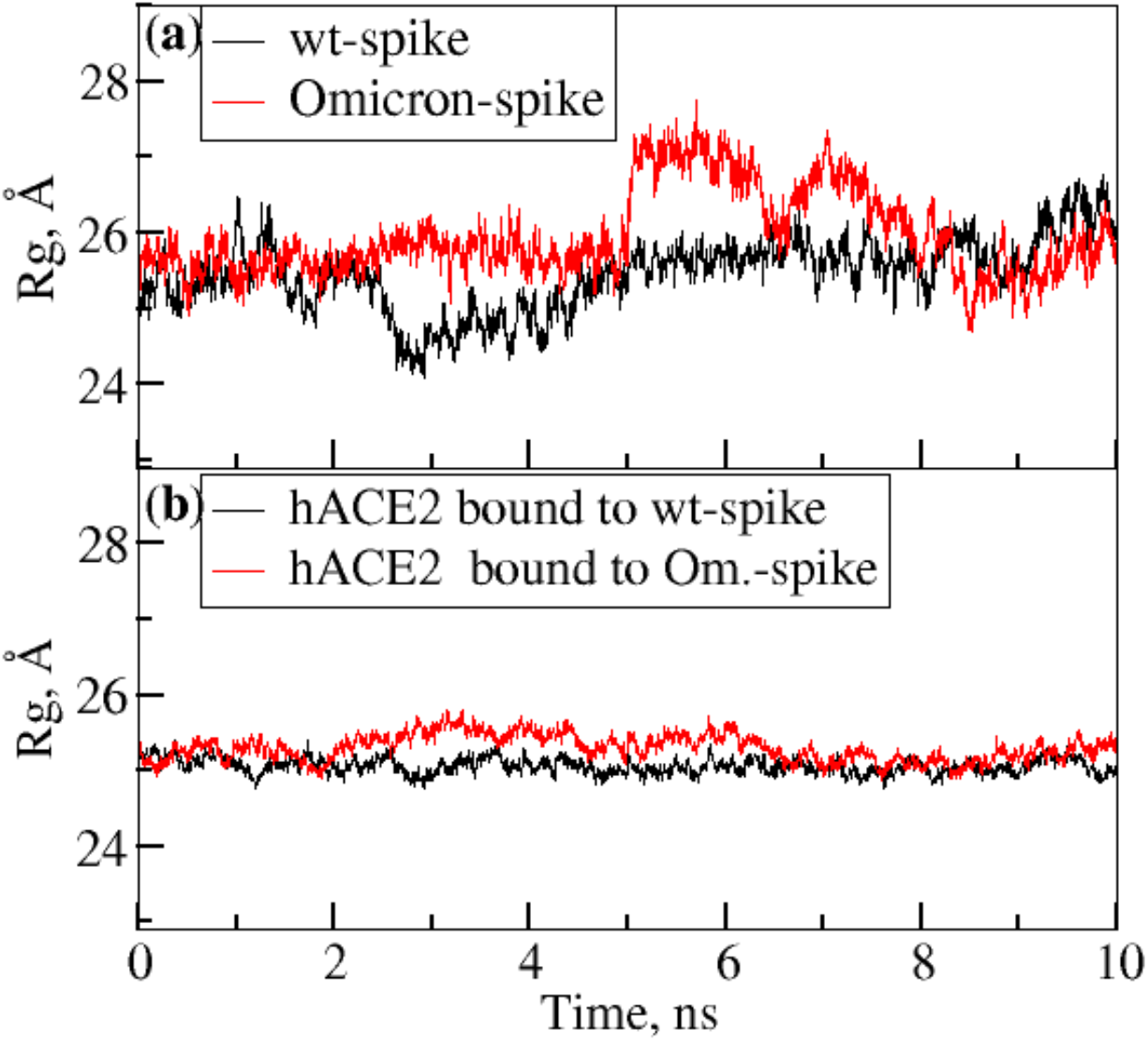
Radius of gyration computed for (a) the spike protein and (b) the hACE2 receptor.

